# Quality Assessment of High-throughput DNA Sequencing Data via Range analysis

**DOI:** 10.1101/101469

**Authors:** M. Oğuzhan Külekci, Ali Fotouhi, Mina Majidi

**Affiliations:** Informatics Institute, Istanbul Technical University, Turkey; Department of Mathematics, University of Tehran, Iran

## Abstract

In the recent literature there appeared a number of studies for the quality assessment of sequencing data. These efforts, to a great extent, focused on reporting the statistical parameters regarding to the distribution of the quality scores and/or the base-calls in a FASTQ file. We investigate another dimension for the quality assessment motivated
with the fact that reads including long intervals having fewer errors improve the performances of the post-processing tools in the down-stream analysis. Thus, the quality assessment procedures proposed in this study aim to analyze the segments on the reads that are above a certain quality. We define an interval of a read to be of desired quality when there are at most *k* quality scores less than or equal to a threshold value *v*, for some *v* and *k* provided by the user. We present the algorithm to detect those ranges and introduce new metrics computed from their lengths. These metrics include the mean values for the *longest, shortest, average, cubic average, and average variation coefficient* of the fragment lengths that are appropriate according to the *v* and *k* input parameters. We provide a new software tool QASDRA for quality assessment of sequencing data via range analysis. QASDRA, implemented in Python, and publicly available at https://github.com/ali-cp/QASDRA.git, creates the quality assessment report of an input FASTQ file according to the user specified *k* and *v* parameters. It also has the capabilities to filter out the reads according to the metrics introduced.

## 1 Introduction

With the spread of high-throughput DNA sequencing, today, not only the research centers, but also the practitioners such as the hospitals, clinics, and even the individuals become customers of the sequencing centers. Each day more sequencing data than the previous is being produced rapidly. This brings a strong necessity to assess the quality of the generated data.

Previous studies [3, 12, 10, 13, 11, 1] for the quality assessment of the DNA sequencing data concentrated on extracting the basic statistical properties such as the mean, median, and standard deviation values of the quality score distribution, where some of those efforts also included the statistical analysis of the base-calls distributions as well, e.g., the `GC` or `N` content ratios.

It is well known that long intervals having fewer errors improve the performances of the post-processing tools in the down-stream analysis of the DNA sequencing data [2]. This brings the idea of evaluating the DNA sequencing data quality via analyzing the lengths of the fragments that are above a certain threshold. Such an assessment requires the explicit definition of *desired–quality* on a read segment.

We propose to identify the quality of fragments by using two parameters *v* and *k*. The v parameter defines a threshold value such that quality scores less than or equal to *v* are assumed to be erroneous. Similarly, in an interval the number of allowed errors, which are defined by *v*, is limited by the parameter *k*. Based on *v*, and *k* parameters, the read segments that include at most *k* scores below *v* are of desired–quality. Finding such ranges has been recently studied in [5] as the inverse range selection queries.

We focus in this study to devise some metrics based on analyzing the lengths of the intervals that include at most *k* quality scores less than or equal to *v* on the quality scores of the reads in an input FASTQ file. The proposed scheme computes a series of metrics for the quality assessment of the input file. We present QASDRA as a new quality assessment tool for DNA sequencing data based on these metrics. QASDRA creates an assessment report that includes the results with various related plots for the input fastq file according to the provided *v*, *k* parameters. Since the fastq files can potentially be so large, random sampling of the reads with a user specified percentage is possible with the QASDRA. Additionally, filtering out the reads that are below the defined threshold is yet another capability of the developed tool.

The outline of the paper is as follows. We briefly review the previous studies in Section 2. Section 3 first introduces the algorithm to answer the inverse range selection queries introduced in [5], and then, describes the proposed metrics along with the reasons that they are devised for. Before the final conclusions, the empirical evaluation on some sample files are given in Section 4.

## 2 Previous Studies

The major tools that have been proposed in the related literature for the DNA sequencing data quality evaluation have focused on statistical distributions of the quality scores, the base–calls, or both. We provide a short review of those tools below.

PIQA [10], was proposed as an extension of the standard Illumina pipeline particularly targeting identification of various technical problems, such as defective files, mistakes in the sample/library preparation and abnormalities in frequencies of sequenced reads. With that purpose it calculates statistics considering the distribution of the A-C-G-T bases. Both the base-calls and their quality scores are considered together.

SolexaQA [3] calculates sequence quality statistics and creates visual representations of data quality for second-generation sequencing data. Default metric is mean quality scores extracted from the reads, but users may also calculate variances, minimum, and maximum quality scores observed. Additionally, the longest read segments with a user defined threshold for minimum quality score is also provided. Based on this calculation, it provides support to trim all reads such that only the longest segment with the user defined threshold remains. The longest fragment detection provided in SolexaQA is a special case of one of our metrics. We discuss this issue in section 3 at the related subsection.

BIGpre [13] provides the statistics such as the distributions of the mean qualities of the reads, and the GC content. The main contribution here has been reported to be the extra features to achieve alignment-free detection of duplicates in the read set.

The quality control and statistics tools in the NGS-QC toolkit [11] is yet another option to retrieve the fundamental statistics of the quality scores and the base-calls. The toolkit includes features to remove low quality reads decided according to the mean quality scores or the base-call distributions.

Similar to SolexaQA, the HTQC [12] performs quality assessment and filtration focusing on statistical distribution of quality scores throughout the input reads with the main motivation of achieving this process faster.

The FastQC software [1] is a commonly used quality control tool. It reports the basic statistics as well as the `GC` or `N` content, per base or per read with a graphical user interface.

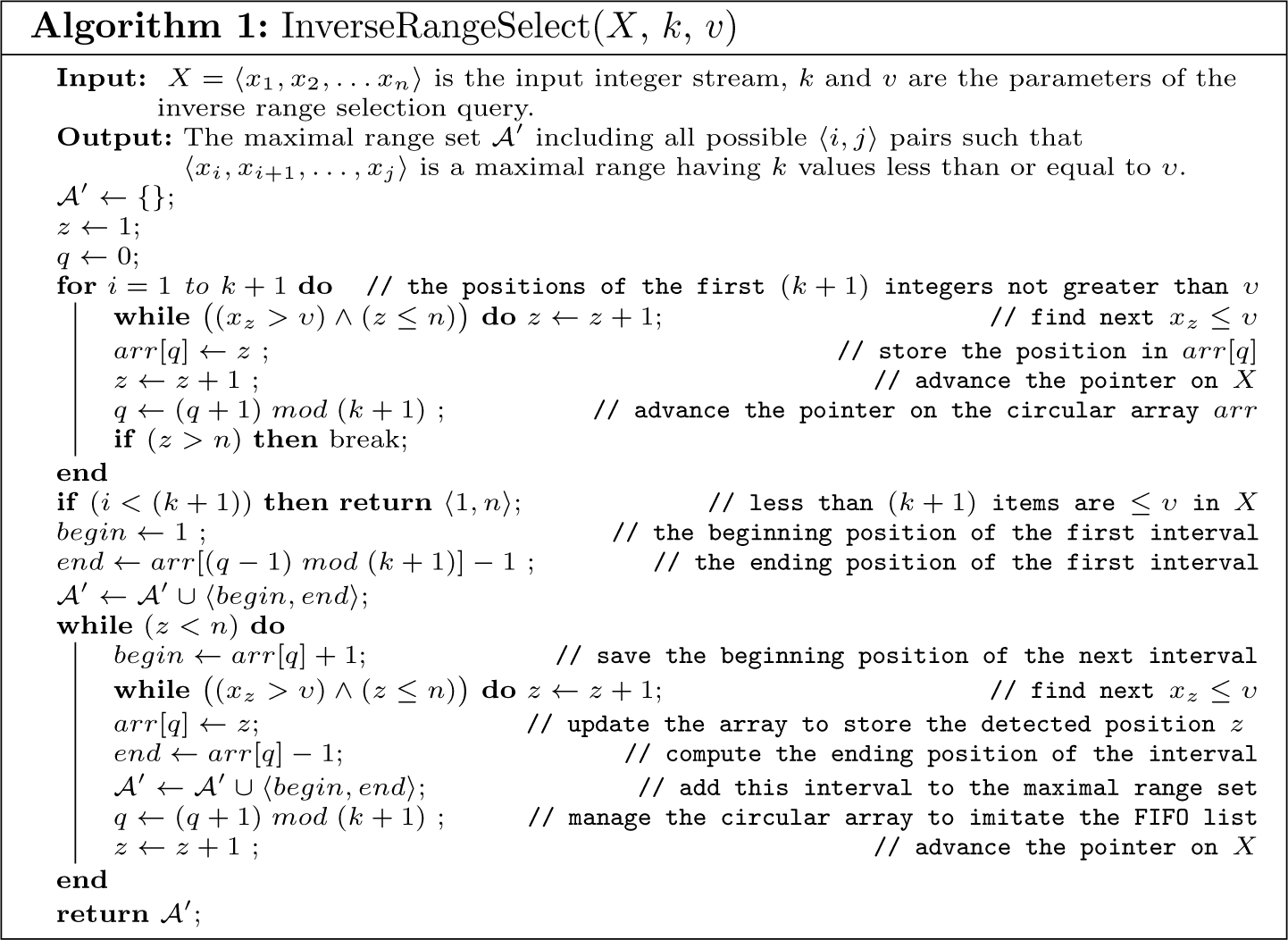

## 3 The Method

The metrics we propose are based on detecting intervals of the reads that contain at most *k* quality scores below a given threshold *v*, which is akin to inverse range selection queries [5].

### 3.1 Inverse Range Selection Queries

On a given integer sequence *X* = 〈*x*_1_, *x*_2_,…*x_n_*〉, the ordinary range selection [4, 6] query *v* ← *R*(*i, j, k*) returns the *k^th^* smallest value *v* in the range (*x_i_*, *x*_*i*+*1*_,…*x_j_*〉, for 1 ≤ *k* ≤ (*j* − *i* + 1) and 1 ≤ *i* ≤ *j* ≤ *n*. Recently the reverse case of this problem has been addressed as the inverse range selection queries [5]. Akin to that, we define in this study the *InvR*(*k,v*) query to return the set 𝓐, which includes all possible 〈*i*, *j*〉 tuples such that in 〈*x_i_*, *x*_*i*+*1*_,…, *x_j_*〉 there are no more than *k* items less than or equal to *v*.

**Definition 1 (Maximal range).** *The range* 〈*x_i_*, *x*_*i*+*1*_,…*x_j_*〉 *denoted by the tuple* 〈*i,j*〉 *in the answer set 𝓐 of the InvR*(*k,v*) *query is a maximal range if there exists no other tuple* 〈*m, n*〉 ∈ 𝓐 *such that m* ≤ *i* ≤ *j* ≤ *n*, which means the 〈*i,j*〉 *interval cannot be expanded either to the right or to the left.*

For example, on *X* = {16, 17, 3, 6, 2,11, 5, 2, 3, 15, 16, 9, 13}, the interval 〈1, 7〉,〈4, 8〉, and 〈6,13〉 are the maximal ranges for the *InvR*(*k* = 2, *v* = 3) query. On the other hand, although 〈2, 7〉 is a valid answer to this query, it is not maximal as it is possible to expand it to the left.

**Lemma 1.** *If* 〈*x_i_*,*x*_*i*+1_,…*x_j_*〉 *is a maximal range, then* [(*x*_*i*−1_ ≤ *v*) ∨ (*i* = 1)] *and* [(*x*_*j*+1_ ≤ *v*) ∨ (*j* = *n*)] *conditions should hold.*

*Proof.* The maximal range 〈*x_i_*,*x*_*i*+1_,…*x_j_*〉 includes *k* items that are less than or equal to *v*, and is not expandable towards right or left. The interval can not be extended to the left when *i* = 1 as this is the leftmost position on *X* that naturally prohibits moving left. In case *i* > 1, if *x*_*i*−1_ > *v*, then 〈*x_i_*,*x*_*i*+1_,…*x_j_*〉 violates the definition of the maximal range, and thus, *x*_*i*−1_ ≤ *v* should hold. In the same way, the expansion to the right is restricted when *j* = *n* or *x*_*j*+1_ ≤ *v*.

*Proof.* There are *k* integers that are less than or equal to *v* in 〈*x_i′_*, *x*_*i′*+1_,…*x_j′_*〉. All of the integers in the intervals 〈*x_i_*,*x*_*i*+1_,…*x*_*i*′−1_〉 and 〈*x*_*j′*+1_, *x*_*j′*+2_,…*x_j_*〉 should be larger than *k*, since there will be otherwise more than *k* items smaller than *v*, which would violate the query. Then, an interval 〈*x_a_*,*x*_*a*+1_,…*x_b_*〉, for *a* ∈ {*i,i* + 1,…, *i′*} and *b* ∈ {*j′*, *j′* + 1,…, *j*}, includes exactly *k* items less than *v*, and holds with the *Q̇*(*v,k*) query.

**Theorem 1.** *The maximal range set of the inverse range selection query InvR*(*k, v*) *on an integer sequence X can be detected in O*(*n*)*-time by using* (*k* +1) *log n bits additional space.*

*Proof.* Following Lemma 1, a maximal range computation requires the knowledge of (*k* + 1) consecutive positions whose corresponding integers are less than or equal to *v* on *X*. A first-in-first-out array *q*[1…(*k* + 1)] keeping the positions of the last observed such (*k* + 1) integers can be maintained while passing over *X* linearly. The size of that array is (*k* + 1)· log *n* bits as each entry is log *n* bits long. After detecting the first such (*k* + 1) integers on *X*, the initial maximal range is 〈*x*_1_…*x*_*q*[*k*+1]−1_〉, which expands from position 1 to the preceding position of the last item in the array. The next maximal range should begin from the succeeding position of the first element in the *q* array. After selecting the starting position of the next maximal range as *q*[1] + 1, the next position on *X* whose corresponding value is less than or equal to *v* is scanned and the *q* array is updated. Since *q* is a FIFO array, this update may be described as shifting all values to the left by one, which disposes *q*[1], and inserting the newly detected position to the end of the array, *q*[*k* + 1]. Now the end of the current maximal range is set to the newly computed (*q*[*k* +1] − 1). Deciding on the start of the next range, updating the array by keep scanning *X* linearly, and setting the end according to the latest update is repeated until all elements in *X* are visited. Since we visit each element of *X* once during the traversal, and maintain an array of size (*k* + 1), the procedure detects all maximal ranges in *O*(*n*)-time and *O*(*k*)-space.

### 3.2 The Metrics

We denote the quality values of a read *t* by *Q*[*t*] = *q*_1_*q*_2_…*q_ℓ_t__*, where *ℓ_t_* is the length of that read. The total number of reads in the input FASTQ file is shown by *N*, and the total length of the reads is *𝓛* = *ℓ*_1_ + *ℓ*_2_ +…+ *ℓ_N_*.

The inverse range selection query *InvR*(*k,v*) on *Q*[*t*] returns the set of maximal ranges as {*r*_1_, *r*_2_,…*r*_*ϕ_t_*_}, where each *r_i_* = 〈*s_i_*,*e_i_*〉 denotes the maximal interval of length |*r_i_*| = *e_i_* − *s_i_* + 1 as *q_s_i__q*_*s*_*i*+1__…*q*_*e_i_*_ such that no more than *k* quality values are less than equal to *v*. The number of detected maximal ranges on *Q*[*t*] is denoted by *ϕ*^*t*^.

The standard variation of the maximal range lengths {*r*_1_, *r*_2_,…*r*_*ϕ_t_*_} is shown by *σ_t_*.

We compute the *InvR*(*k,v*) on each read based on the parameters *k* and *v* provided by the user, and then, calculate the following per read based on the detected maximal range lengths (MRL).

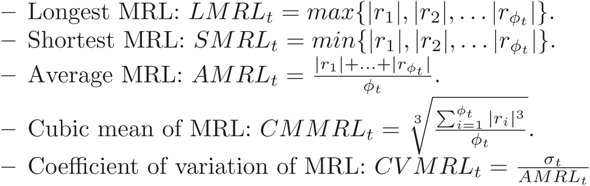

The proposed quality assessment metrics based on the individual values calculated per read are defined below.

Average *longest* maximal range length (ALMRL)

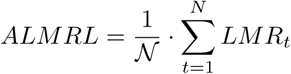

The performance of the downstream processing on DNA sequences increase with longer intervals having less errors. For instance, it had been shown in [3] that filtering the low quality segments of the reads improve the assembly performance. Thus, ALMRL aims to evaluate the quality by measuring the lengths of the segments that are defined to be of enough quality by the inverse range selection query.

Notice that this metric is akin to the dynamic trimming of the SolexaQA [3] that detects the longest read segment, where the minimum quality value is above a threshold. The process regarding to that in SolexaQA is a special case of ALMR metric by setting the *k* value to 1, where the proposed tool allows variable number of bases instead of one to be below the threshold value. This extension make sense when one uses methods that can handle multiple errors on the reads. For example, the alignment applications such as the BWA [8], Bowtie [7], and others[9] have mechanisms to handle more than one error efficiently.

Larger ALMRL scores indicate better quality. Since the perfect LMRL value of a read is its length, which indicates the number of quality scores below v are at most *k* throughout the read, the best ALMRL score of a fastq file is actually its average read length.

Average *shortest* maximal range length (ASMRL)

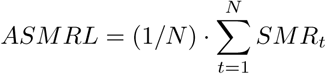

The shortest maximal range on a read indicates the smallest distance in which there are *k* quality scores that are below *v*. This value becomes *k* in the worst case, where all the low quality values appear subsequently. When the *SMRL^t^* value of a read is significantly small, it means there is a burst error, where the erroneous values appear very close to each other.

Thus, the ASMRL metric might be useful for the purpose of measuring the distribution uniformity of the low quality positions. For instance, when those erroneous positions do not appear close, larger ASMRL values are expected.

Grand average of maximal range lengths (GAMRL)

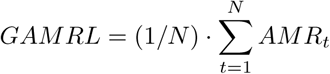

There might be, and most possibly will be, more than one maximal range on a read. The mean value of those maximal range lengths are computed per each, and the grand average of those means can be used to evaluate the overall performance of the sequencing process assuming that higher values of GAMRL indicates better quality. It is expected that on a given read there appears a segment of length GAMRL withholding the queried *v*, *k* criteria. However, this measure is a bit coarse, and due to that, we introduce additional metrics below to support more detailed analysis of the detected segments.

Average cubic means of maximal range lengths (ACMRL)

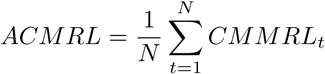

When the maximal range lengths are detected on a read, we would like to devote more weight to the longer ones than the shorts. With this aim, despite the GAMRL measure, we also compute the cubic mean of the MRLs on a read. Notice that cubic mean, which is a generalized mean computation 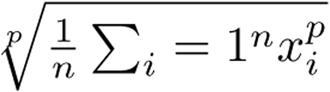 with p=3, favors the long MRL values more. For instance assume the detected MRLs on two different reads are 〈20, 30,40〉 〈30, 30, 30〉. Although their averages are both 30, their cubic means are 32.07 and 20.80. Here we can see the difference made by longer segments which shows the power of longer reads.

The ASMRL metric mainly evaluate the fragmentation and burst errors in the reads. However, even in case of high fragmentation and burst errors, there might still be enough long segments that can be helpful in downstream analysis. The ACMRL metric aims to provide a way of measuring the longest maximal range lengths associated with the number of maximal ranges detected. We would like to increase the power of the longer maximal ranges, and thus, tried generalized means with different values, where empirically decided on cubic mean as the best value to measure this.

Average of coefficient of variations of MRL (ACVMRL)

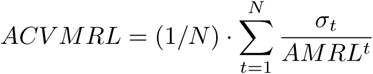

The coefficient variation (CV) is defined as the ratio of the standard deviation *σ* to the mean *μ*. The coefficient variation is useful because the actual value of CV is independent of the unit in which the measurement has been taken, so it is a dimensionless number. For comparison between data sets with different units or widely different means, one should use the coefficient of variation instead of the standard deviation.

For example, a data set of [100,100,100] has constant values its standard deviation is 0 and its average is 100 so *C_v_* =0, a data set of [90,100,110] has more variability. its standard deviation is 8.165 and its average is 100 so *C_v_* =0.08165, and a data set of [1,5,6,8,10,40,65,88] has more variability again. its standard deviation is 30.78 and its average is 27.875 so *C_v_* = 1.104. Since each read has different number of maximal ranges lengths with different means, this concept illustrates the coherence of the data in terms of MRLs computed, where higher values indicate less uniformity among the maximal range lengths. Thus, having a small ACVMRL is good in terms of quality, and indicates that one may expect to be more confident to observe the computed *average* values in a randomly selected read.

## 4 Empirical Evaluation

We present the results of our assessment scheme first on the data sets used by the FastQC [1] program evaluation. This data includes two (old) FASTQ files generated by the Illumina equipment, which are examples of good and bad sequencing. Roughly there are 250K reads in good data set, and 325K in the bad data set. The reads are 40 bases long.

**Fig.1.**
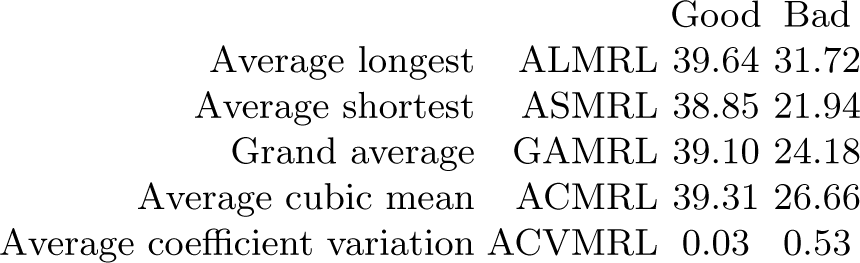
The quality assessment of the good and bad Illumina sequencing data from FastQC [1] with the proposed metrics with *k* = 2 and *v* = 20.

**Fig. 2.**
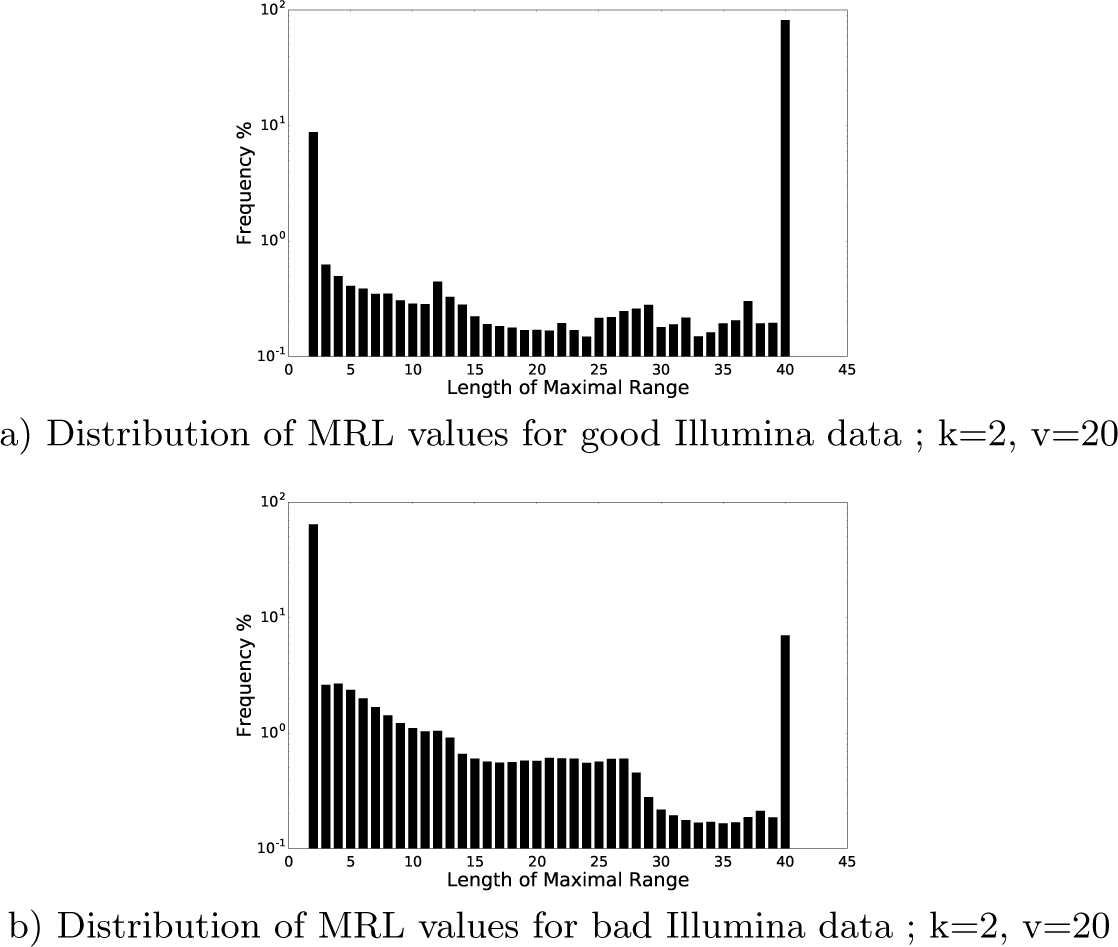
Distribution of maximal range lengths on good and bad Illumina data for *k* = 2, *v* = 20.

An expected difference in the ALMRL of the data sets have appeared such that the average length of the longest maximal range is 39 in good data and 31 in bad data. Similarly the ASMRL as 21.94 on bad data versus 38.85 on good data reflects the quality difference.

In terms of GAMRL (39.10 versus 21.18) and ACMRL (39.31 versus 26.66) values, we observe that the difference in GAMRL is sharper. That reminds us although the mean MRL in bad data is not that much good, there still appears some long intervals in the reads so that the ACMRL value improves.

Notice that the ACVMRL values for good and bad data are quite distinct, where it is much larger on bad data set. Due to that high value, we can think that the diversity of the maximal ranges in bad data is quite high, where the distribution is much smooth in good data. This can also be partially observed on Figure 4. The full reports for the good and bad data are provided in the appendices. We also present the results of our study on more recent files generated by the different sequencing platforms with the Illumina, IonTorrent, and PacBio equipments. we have used individual NA12878 as published by the Coriell Cell Repository. The data files used are ERR091571_1.fastq and ERR091571_1.fastq concatenated to one file ERR091571.fastq, High coverage reads for NA12878, from Illumina, SRR1238539.fastq for IonTorrent, including 183976176 reads of lengths varying between 25 and 396, chemistry_3 for PacBio, consisted of 8 fastq files concatenated to one, including 654547 reads of lengths varying between 50 and 33230.

The results of the quality assessment with the proposed technique are given in Figure 3, and the distributions of the MRLs per each platform are depicted in Figure 4.

**Fig. 3.**
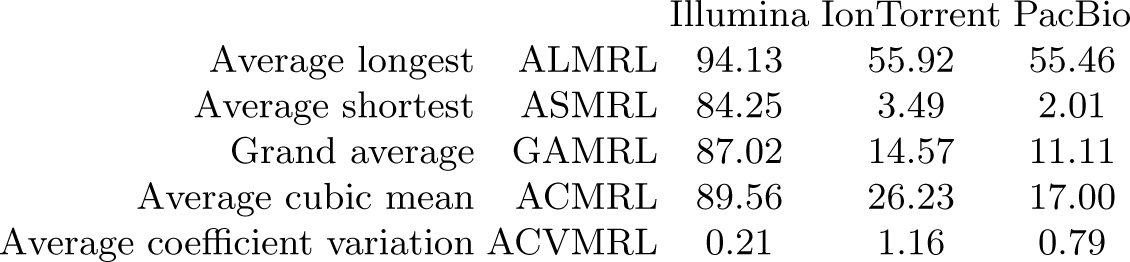
The quality assessment of the selected fastq files generated by the Illumina, IonTorrent and PacBio platforms.

**Fig. 4.**
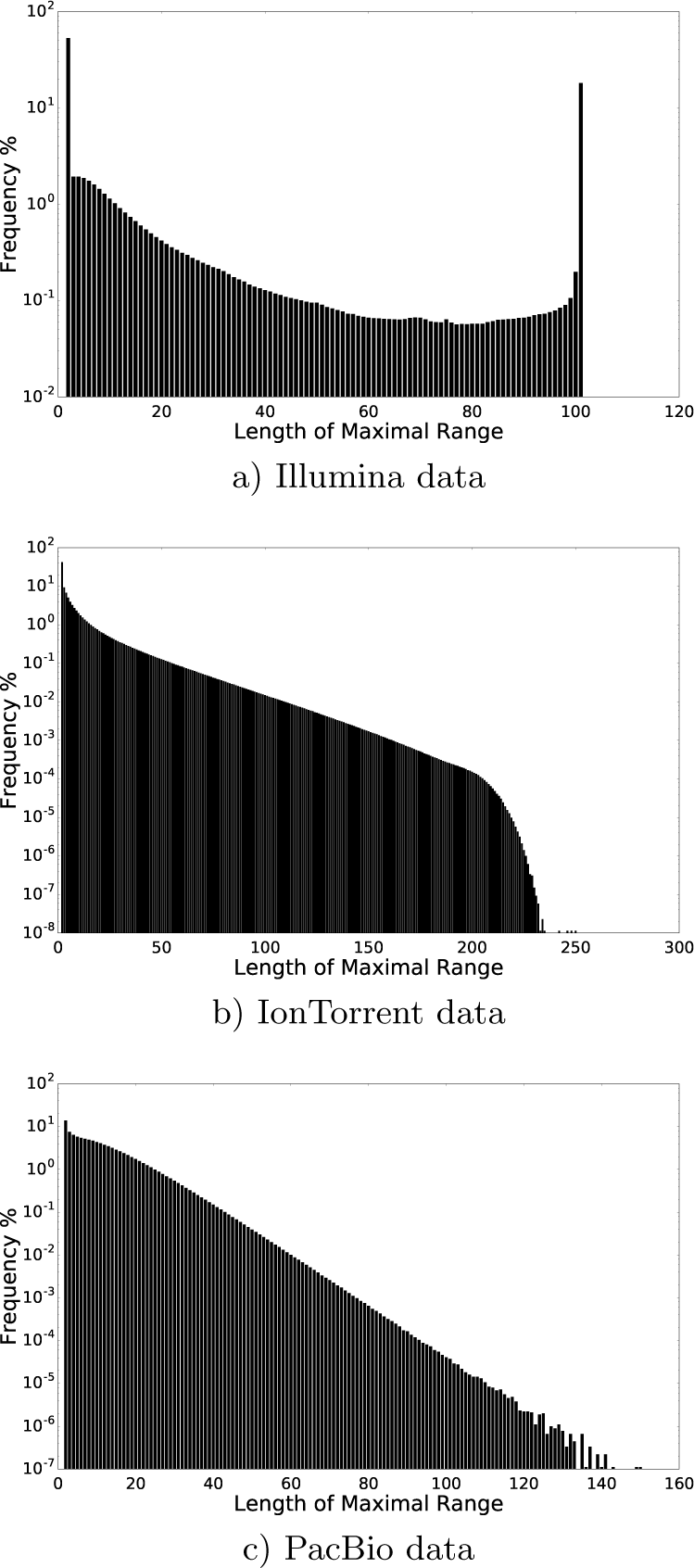
Distributons of the MRL on the selected data files of Illumna, IonTorrent, and PacBio data copmuted with *k* = 2, and *v* = 20. Notice that *v* = 7 is used on PacBio data that approximately corresponds to *v* = 20 on other platforms.

We observed that the Illumina reads, according to the given parameters *k* = 2 and *v* = 20, include longer maximal ranges according to the ALMRL, ASMRL, and GAMRL metrics. On these measures, the IonTorrent and PacBio platforms returned similar results particularly on ALMRL, which means the average longestTorrent and 2.01 for the PacBio data. This reminds us that on the selected data sets, whenever a quality score below 20 is observed, its very near neighbors are also usually below that quality, and hence, the ASMRL values are that much small. Considering the GAMRL metric, the tested IonTorrent data provides slighly longer contigious blocks of desired-quality. The average cubic mean being 26.23 for the IonTorrent and 17.00 for the PacBio indicates that although the ALMRL, ASMRL, and GAMRL values are close, the reads in the IonTorrent data include more longer intervals than the PacBio. However, the high value of ACVMRL in the IonTorrent shows that the PacBio data is more uniformly distributed. In the ACVMRL metric, the Illumina shows a much nicer distribution.

The computed metrics are highly sensitive to the selected *k* and *v* parameters. Thus, we have included quality assessment experiments repeated on the same data with different parameters. The results are shown in Table 1. Notice that on PacBio and IonTorrent platforms, the shortest maximal range values are quite small, which means in general the low-quality base-calls appear very close. The larger ASMRL value on Illumina data shows that the low-quality positions are more uniformly spread here.

**Table 1.** Quality assessment of some selected fastq files with different *v* and *k* values.

| File Info | $k$ Value | $v$ Value | ALMRL | ASMRL | GAMRL | ACMRL | ACVMRL |
| --- | --- | --- | --- | --- | --- | --- | --- |
| Platform: Illumina<br>Sequence Data: NA12878<br>Name: ERR091571.fastq<br>Number of Reads: 422875838<br>Quality Scores: (2,41)<br>Read Lengths: (101,101) | 2 | 20 | 94.13 | 84.25 | 87.02 | 89.56 | 0.21 |
|  |  | 30 | 83.86 | 62.87 | 69.25 | 74.47 | 0.41 |
|  | 3 | 20 | 95.10 | 86.64 | 88.94 | 90.99 | 0.16 |
|  |  | 30 | 86.03 | 68.29 | 73.60 | 77.73 | 0.31 |
|  | 4 | 20 | 95.79 | 88.33 | 90.31 | 92.04 | 0.13 |
|  |  | 30 | 87.59 | 72.09 | 76.70 | 80.09 | 0.24 |
| Platform: ION Torrent<br>Sequence Data: NA12878<br>Name: SRR1238539.fastq<br>Number of Reads: 183976176<br>Quality Scores: (3,38)<br>Read Lengths: (25,396) | 2 | 20 | 55.92 | 3.49 | 14.57 | 26.23 | 1.16 |
|  |  | 30 | 3.49 | 2.00 | 2.00 | 2.11 | 0.12 |
|  | 3 | 20 | 63.18 | 5.84 | 19.66 | 31.74 | 0.98 |
|  |  | 30 | 4.54 | 3.00 | 3.00 | 3.12 | 0.10 |
|  | 4 | 20 | 69.52 | 8.58 | 24.66 | 36.89 | 0.86 |
|  |  | 30 | 5.58 | 4.00 | 4.00 | 4.14 | 0.08 |
| Platform: PacBio<br>Sequence Data: NA12878<br>Name: chemistry_3.fastq<br>Number of Reads: 654547<br>Quality Scores: (0,14)<br>Read Lengths: (50,33230) | 2 | 7 | 55.46 | 2.01 | 11.11 | 17.00 | 0.79 |
|  |  | 11 | 22.14 | 2.00 | 4.13 | 6.38 | 0.74 |
|  | 3 | 7 | 63.84 | 3.05 | 15.32 | 21.54 | 0.67 |
|  |  | 11 | 25.70 | 3.00 | 6.00 | 8.32 | 0.60 |
|  | 4 | 7 | 71.73 | 4.13 | 19.52 | 26.04 | 0.61 |
|  |  | 11 | 29.07 | 4.00 | 7.87 | 10.25 | 0.52 |
| Platform: Illumina<br>Sequence Data: Exome sequencing of PBCF cell line<br>Name: NCI-PBCF-HTB68.pson0001.Exome.fq<br>Number of Reads: 46192166<br>Quality Scores: (2,41)<br>Read Lengths: (100,101) | 2 | 20 | 92.87 | 84.30 | 86.92 | 88.91 | 0.17 |
|  |  | 30 | 80.97 | 61.86 | 67.98 | 72.40 | 0.37 |
|  | 3 | 20 | 94.07 | 86.72 | 88.91 | 90.52 | 0.13 |
|  |  | 30 | 83.52 | 67.17 | 72.39 | 75.88 | 0.28 |
|  | 4 | 20 | 94.92 | 88.44 | 90.33 | 91.68 | 0.10 |
|  |  | 30 | 85.37 | 71.01 | 75.59 | 78.44 | 0.22 |
| Platform: ION Torrent<br>Sequence Data: NA12878/ion_exome<br>Name: bb17523_PSP4_BC20.fastq<br>Number of Reads: 40008921<br>Quality Scores: (3,38)<br>Read Lengths: (12,347) | 2 | 20 | 71.81 | 7.57 | 24.65 | 38.61 | 1.03 |
|  |  | 30 | 3.84 | 2.00 | 2.00 | 2.15 | 0.16 |
|  | 3 | 20 | 81.10 | 12.53 | 32.68 | 46.68 | 0.87 |
|  |  | 30 | 4.91 | 3.00 | 3.00 | 3.17 | 0.12 |
|  | 4 | 20 | 89.13 | 18.05 | 40.38 | 54.15 | 0.76 |
|  |  | 30 | 5.98 | 4.00 | 4.00 | 4.19 | 0.10 |
| Platform: PacBio<br>Sequence Data: CHM1 subreads<br>Name: m130929_024849_42213.fastq<br>Number of Reads: 72708<br>Quality Scores: (0,14)<br>Read Lengths: (35,35488) | 2 | 7 | 52.92 | 2.01 | 10.66 | 16.17 | 0.76 |
|  |  | 11 | 21.33 | 2.00 | 3.98 | 6.13 | 0.73 |
|  | 3 | 7 | 60.83 | 3.03 | 14.72 | 20.49 | 0.65 |
|  |  | 11 | 24.76 | 3.00 | 5.81 | 8.00 | 0.58 |
|  | 4 | 7 | 68.30 | 4.09 | 18.78 | 24.78 | 0.58 |
|  |  | 11 | 27.98 | 4.00 | 7.64 | 9.88 | 0.51 |

## 5 Conclusion

The statistical properties of the distributions regarding both the quality scores and the base–calls of a sequencing experiment have been extensively explored in previous studies. We have presented an alternative approach to the quality assessment of sequencing data by analyzing the maximal ranges, which are defined as the longest segments in which no more than *k* scores are less than or equal to *v*. The software developed with Python for the proposed metric is available at https://github.com/ali-cp/QASDRA.git for public use.

The sequencing centers or the consumers of those centers can use the tool to evaluate or benchmark their data. In the near future, it might be necessary to define the international standards of good sequencing data, where we believe the approach presented in this study might help in creating such standards. The metrics introduced in this study may serve for clustering/classifying the reads from the different platforms, or for the overall successes of the sequencing centers.

## Acknowledgements

We thank S. Andrews from the Babraham Bioinformatics for providing us the good and bad Illumina data sets they reported in their FastQC software.

This work has been partially supported by the TUBITAK-114E293 research grant of Turkey.

## 6 Appendices

### QASDRA - Quality Assessment of Sequencing Data via Range Analysis

File Name: gooddata.fastq / Date: Mon Jan 16 12:53:26 2017

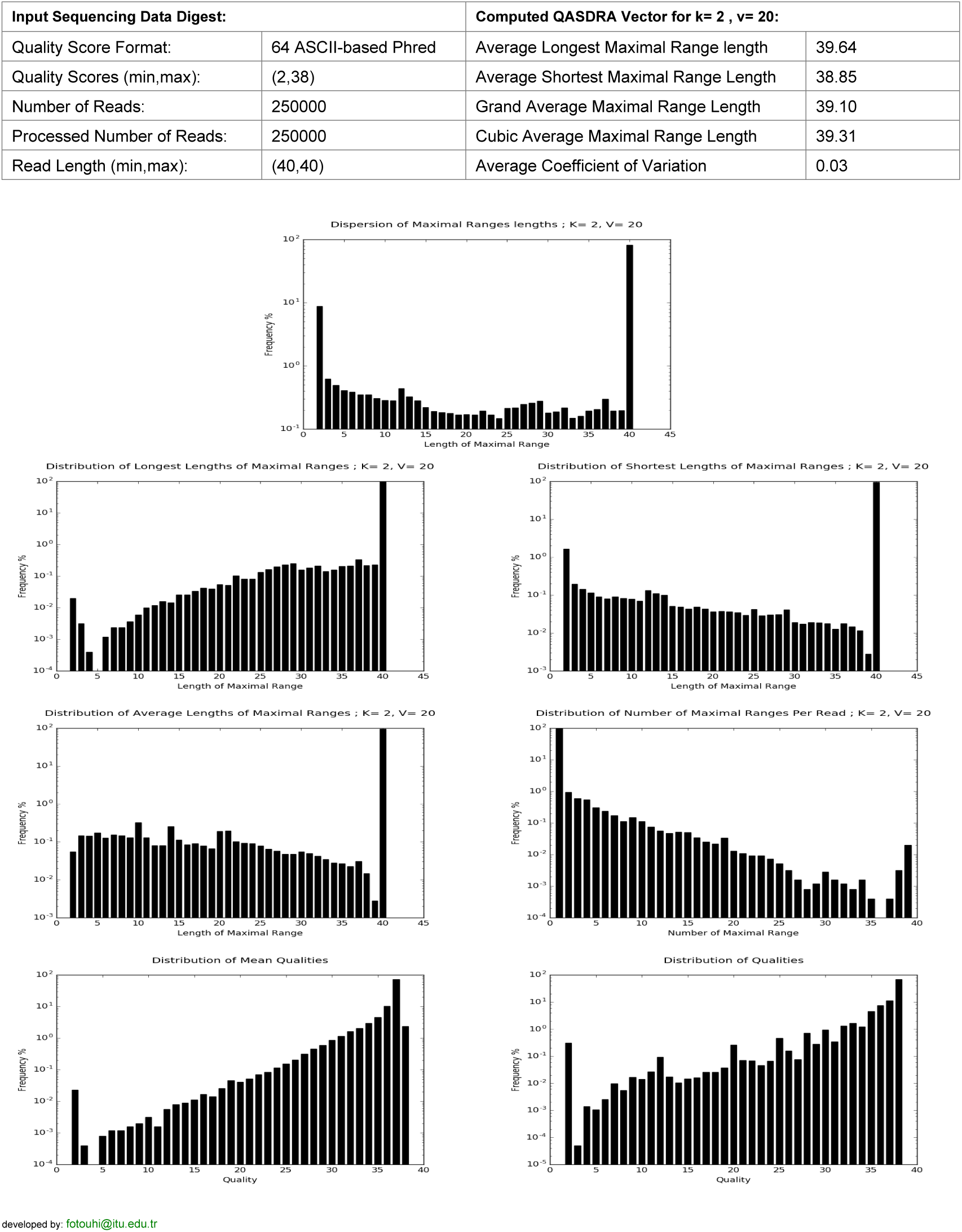

### QASDRA - Quality Assessment of Sequencing Data via Range Analysis

File Name: baddata.fastq / Date: Mon Jan 16 12:54:09 2017

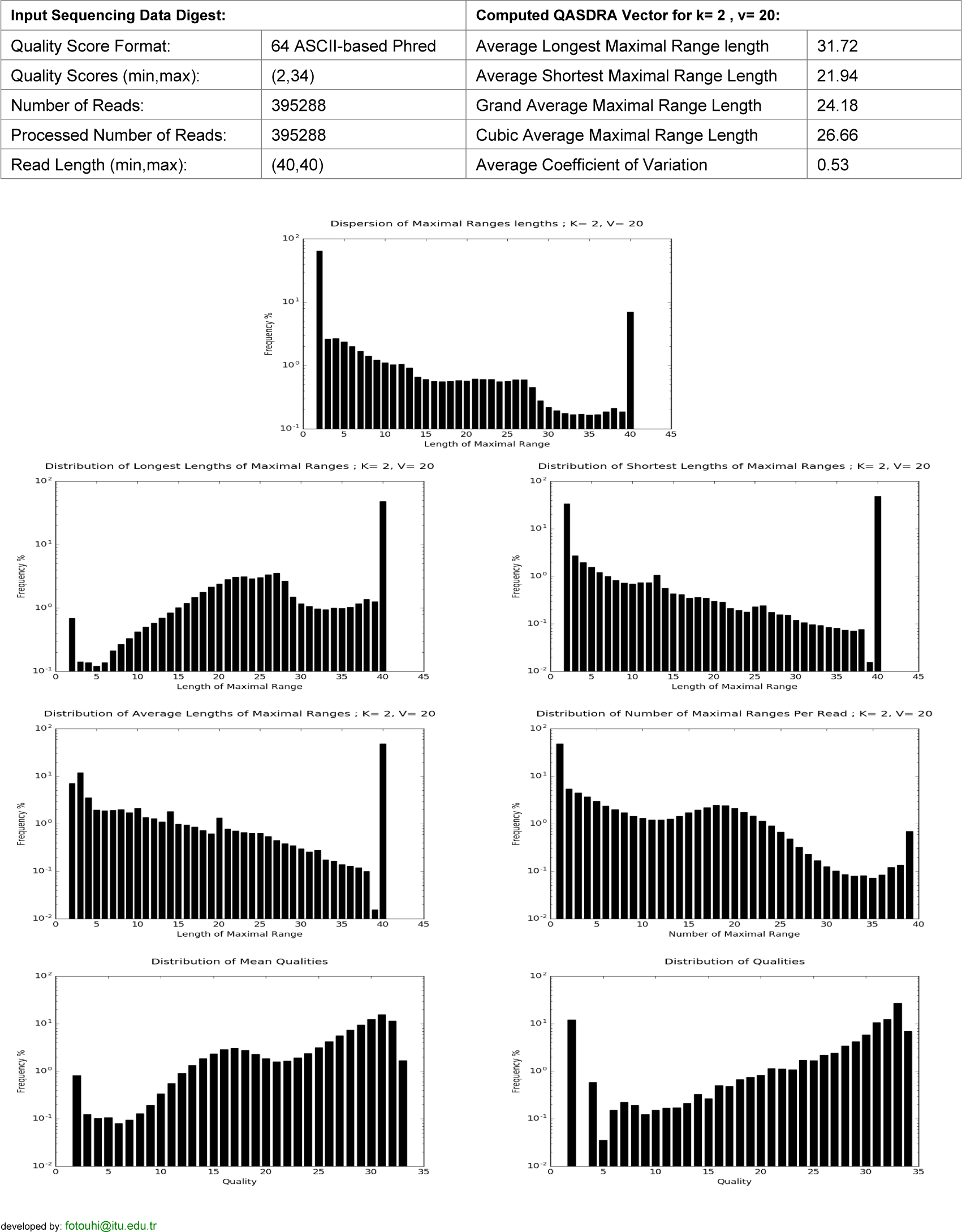

